# Smart computational exploration of stochastic gene regulatory network models using human-in-the-loop semi-supervised learning

**DOI:** 10.1101/490623

**Authors:** Fredrik Wrede, Andreas Hellander

## Abstract

Discrete stochastic models of gene regulatory network models are indispensable tools for biological inquiry since they allow the modeler to predict how molecular interactions give rise to nonlinear system output. Model exploration with the objective of generating qualitative hypotheses about the workings of a pathway is usually the first step in the modeling process. It involves simulating the gene network model under a very large range of conditions, due to the the large uncertainty in interactions and kinetic parameters. This makes model exploration highly computational demanding. Furthermore, with no prior information about the model behavior, labour-intensive manual inspection of very large amounts of simulation results becomes necessary. This limits systematic computational exploration to simplistic models. We address this by developing an interactive, smart workflow for model exploration based on semi-supervised learning and human-in-the-loop labeling of data. The proposed methodology lets a modeler rapidly discover ranges of interesting behaviors predicted by a model. Utilizing the notion that similar simulation output is in proximity of each other in feature space, the modeler can focus on informing the system about what behaviors are more interesting than others instead of configuring and analyzing simulation results. This large reduction in time-consuming manual work by the modeler early in a modeling project can substantially reduce the time needed to go from an initial model to testable predictions and downstream analysis.

## Introduction

A substantial level of single-cell variability both between cell-lines but also between individual cells in the same cell-line has long been a prediction of both stochastic simulations in systems biology ^1–6^ and experimental observations^7, 8^. Single-cell transcriptomics such as scRNA-Seq^9^ and single-cell proteomics^10^ have offered snapshots of this variability in RNA and protein expression for a large number of genes. Comparative studies overlaying such transcriptomics and proteomics data have also recently highlighted how protein expression may or may not be correlated with the expression of its mRNA both in E. Coli^11^ and recently in mouse embryonic stem cells^10^, hinting to the complexity in regulation arising from non-linearly interacting pathways. While single-cell omics and modern analysis pipelines make it possible to quantify differentially expressed genes across a population^12^, these techniques do not enable direct study of the precise mechanisms that give rise to the observed variability. Quantitative, stochastic and dynamic models of these gene regulatory networks (GRN), on the other hand, can be used to predict how such nonlinear interactions result in observed gene expression profiles and how they can explain cell-cell variability. Discrete stochastic simulation of the spatial and temporal dynamics of intracellular biochemical reaction networks has thus become a central tool to study GRNs in system biology^2, 5, 6, 13^, ^14^. These simulation methods capture intrinsic noise due to low copy numbers of the involved molecular species^15^, and can be used to make predictions about both the system dynamics and the variability in behaviors across a cell population.

However, the use of modeling to generate predictions in the absence of prior knowledge about system behavior is challenging due to the large uncertainty in the reaction network interactions and the large amounts of unknown kinetic rate parameters. To explore models under this uncertainty with the objective of obtaining a qualitative understanding of the type of interesting behavior the model suggest about the underlying system, the modeler uses parameter sweeps in which the parameters are varied over a large range and the simulation results are analyzed in postprocessing steps. There are two main problems with this approach. First, in order to encode the critera for postprocessing analysis, the modeler needs to have prior hypotheses about model behavior. Thus there is a substantial risk to miss interesting dynamics that was not already anticipated. Second, due to the curse of dimensionality global parameter sweeps become very large and will require massive computations and time-consuming manual work by the modeler to inspect output time series data and manually fine-tune sampling of the parameter space. In order to address both these problems, we have developed tools based on human-in-the-loop semi-supervised machine learning with the objective to greatly reduce both the manual work involved in going from an initial model to insight about the principle behavior of the model.

We propose a smart workflow framework that utilizes cycles of simulation, feature generation, semi-supervised learning and human-in-the loop labeling. Here we use semi-supervised to infer different qualitative system behaviors of known realizations of the model based on feedback from the modeller. This feedback is in the form of letting the modeller label qualitative behaviors or interesting realizations. To enable this we need ways to visualize and interpret large collections of simulated results. We transform individual molecular trajectories into summary statistics, or features, and visualize each individual realization in a reduced dimensional space. By using interactive scatter plots where each point corresponds to a realization with parameter values, trajectories and summary statistics, we enable the modeller to much more easily explore global parameter sweeps, since the localization of points provides information about the similarity between them. By interactively inspecting (clicking) on realizations in the reduced space, the modeller will be able to quickly inspect the corresponding trajectories for a particular molecular species. The modeller then label only a few realization according to her interpretation of the behaviors and we then use specialized semi-supervised models to infer labels of unlabeled realizations. In doing so, we ask the modeller for feedback on classification of realizations with high ambiguity. This process is referred to as active learning. This allows us both to improves the semi-supervised model, and to present data points of possibly unseen qualitative behaviors to the user.

The main contribution of our smart workflow tools is to greatly reduce the amount of manual work required by the modeler by 1. Automating the process of generating feature (e.g which can be used to engineer summary statistics) in the absence of prior information, and 2. Replacing model-specific manual postprocessing with large-scale autonomous analysis enabled by user interactivity. In addition, the methodology enables more efficient refined analysis in later modeling cycles. By exploitatively selecting (filtering) different behaviors seen in the high-throughput realization data, modelers can easily choose to neglect data that is not considered interesting. This has the potential to drastically reduce storage requirements by avoiding storing non-interesting realizations. Moreover, the information about a modeler’s preferences can be used to train classifiers and use them to guide the sampling process of the parameter space where the exploration of “interesting” regions are accelerated. This way of training classifiers based on modeler input can be seen as a way to engineer objective functions that can be used in systematic downstream sampling algorithms that require prior information.

## Results

### Smart exploration of a model of a regulatory positive-negative feedback loop maps out interesting behavior and engineers features with minimal manual work

We demonstrate the methodology using a GRN model of a circadian clock based on positive and negative regulatory elements. The model, which has 9 species, 18 reactions and 15 reaction rate parameters involves two genes with the translated transcription factors acting as positive and negative regulatory elements, respectively. This model was developed in one of the first studies highlighting the importance of intrinsic molecular noise in systems biology. The model can give rise to robust oscillations in the presence of noise, and the parameter values and the initial condition to achieve robust oscillations with a period of 24 hours is well known^14^. However, for demonstration purposes, here we assume no *a priori* knowledge about the system dynamics and demonstrate how our interactive workflow lets us efficiently explore the high-dimensional parameter space, discover oscillatory dynamics as the principal behavior and interactively label such behavior as interesting. The exploration process for this model example can be divided into three logical phases; I. the initial exploration and labeling, II.label propagation and III. zooming in on region of interest (ROI) and engineer summary statistics.

#### Phase 1: An initial exploration reveals oscillations as a principal behavior of the model

Figure 1 illustrates the first initial phase of the smart model exploration workflow. Using Jupyter interactive notebooks, the modeller has a central role to guide the workflow during exploration. Assuming that the model contains high diversity in the model behavior, a global sampling of parameter space, with an initial small batch size, can ease the process of identifying interesting behaviors for the modeller by keeping the visual information at a comprehensible level. Here we chose an initial batch size of 1000 realizations which span the entire initial sweep design (Supplementary information Table 1), which is provided by the modeller. Further, each parameter point is simulated in parallel with the stochastic simulation algorithm (SSA). A natural bottleneck in the simulator when using a wide span of parameter values is the possibility of parameter combinations that leads to uncontrolled growth, or explosion, of some chemical species. This will cause the simulation time to become very large. To avoid this we set a timeout (50s/realization) for the simulation, effectively filtering our bio-physically unrealistic, exploded trajectories. Once the simulation batch has completed, each simulated trajectory of the species passes through a summary statistics, or feature, generation process using TSFRESH. A minimal set of summary statistics comprising of extreme values, mean, median, standard deviation, variance and the sum are initially generated, converting the raw time series trajectory into an array of features. To visualize and interact with the data, a dimensional reduction (DR) of the feature space corresponding to a particular species is performed. In this example we use UMAP^16^ but other options are possible, such as t-SNE^17^ and PCA. At this point, the modeller is able to interact with each individual parameter point in a low dimensional representation of the feature space. By clicking on a parameter point, the corresponding trajectory of a specified species is shown and points in close proximity to each other have similar behavior, which aids the modeler in initial exploration (see Supplementary video I). Observe that the modeller has to tune and play with the hyperparameters of the DR method as part of the exploration. Figure 1 shows the reduced feature space on the Activator protein using UMAP. In our example, the first batch resulted in 244 out of 1000 realizations completing within the threshold used (50s/realization), the remainder resulting in unphysical explosion of one or more species. As the modeller explore the data she will be able to label interesting clusters or individual parameter points according to her preference. Here, we observed that the left hand side of the reduced feature space corresponds to trajectories with consistent low copy numbers of the transcription factors (Figure 1 A), and they are labeled as non-interesting. The upper right cluster on the other hand have more reasonable copy numbers and a bursting behavior of high frequency (Figure 1 B), a phenomena which can (with parameter tweaking) result in oscillations. As the cluster elongates to the right of the feature space, the higher the copy number becomes. With our preferences these were labeled as interesting. Further, we observe two outlier clusters, where the labeling become more uncertain. One parameter point from each cluster was labeled. (Figure 1 C). Phase 1 of the workflow can be repeated as many times as desired to add more data and support for individual classes. We conducted three batches of the same size as the first batch and following the same uniform sampling of the parameter space. A few more samples was added to the interesting and non-interesting class, respectively.

**Figure 1.**
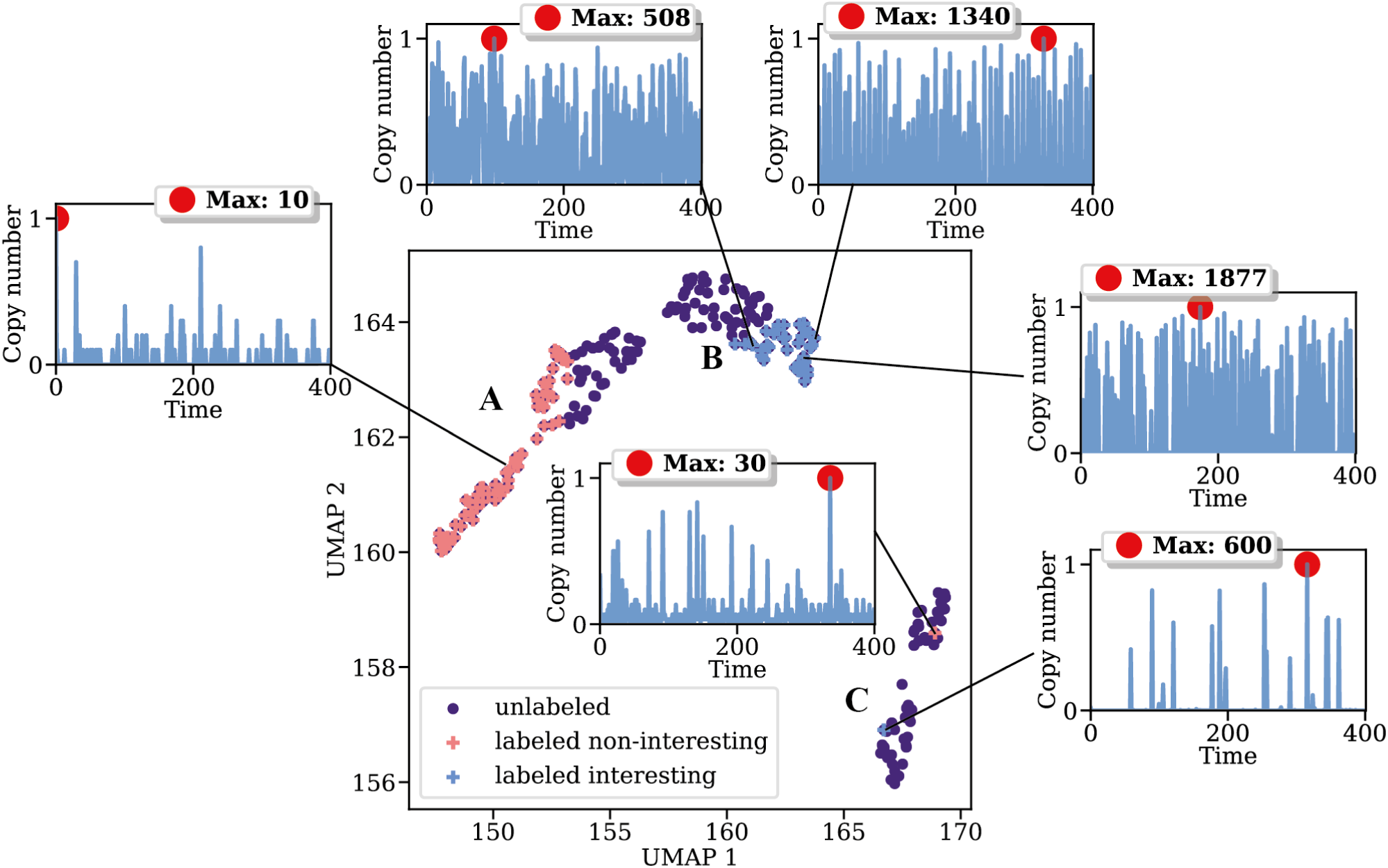
A first phase of the smart model exploration workflow. After an initial, coarse parameter sampling, and after generating summary statistics using a minimal set of time series features, the simulation output (here represented in feature space of the Activator protein) is embedded into a lower dimensional representation using UMAP. This plot is then presented to the modeler as an interactive plot in a Jupyter Notebook and the modeler quickly inspects a few representative samples in the data clusters. The modeler will be able to label samples according to their preferences.

#### Phase 2: Semi-supervised learning to propagate labels to unlabeled time series

Phase 1 results in the modeler gaining an initial understanding of the model’s behavioral diversity, and in some few initial labeled parameter points representing this diversity. After simulating more data and performing a new dimension reduction on the data, the workflow uses a semi-supervised learning algorithm to predict labels of the unlabeled realizations. Here, we use label propagation in the reduced dimensional space, which uses the properties of Gaussian Random Fields and harmonic functions^18^. After propagating the labels, every single parameter point in the current state of the exploration will have an associated probability of belonging to a label (Figure 2). Since label propagation uses the notion that similar data points have similar labels, the modeller can use this information to “guide the eye” in an otherwise very cluttered data set, which can markedly reduce manual work. Further, the modeller will be able to map out points with high label uncertainty by directly observing the probability (green color in Figure 2). Usually the highest uncertainty is at the boundary of two labels in feature space or in the outliers. One way of measuring the uncertainty or information gain associated with groups or individual realizations can for example be entropy-based or based on the generalization error^19^. By letting the modeller label a few of the realization with the highest e.g entropy, we will efficiently gain more information about the preferred label distributions and thus get a better semi-supervised model. This is commonly referred as active learning.

**Figure 2.**
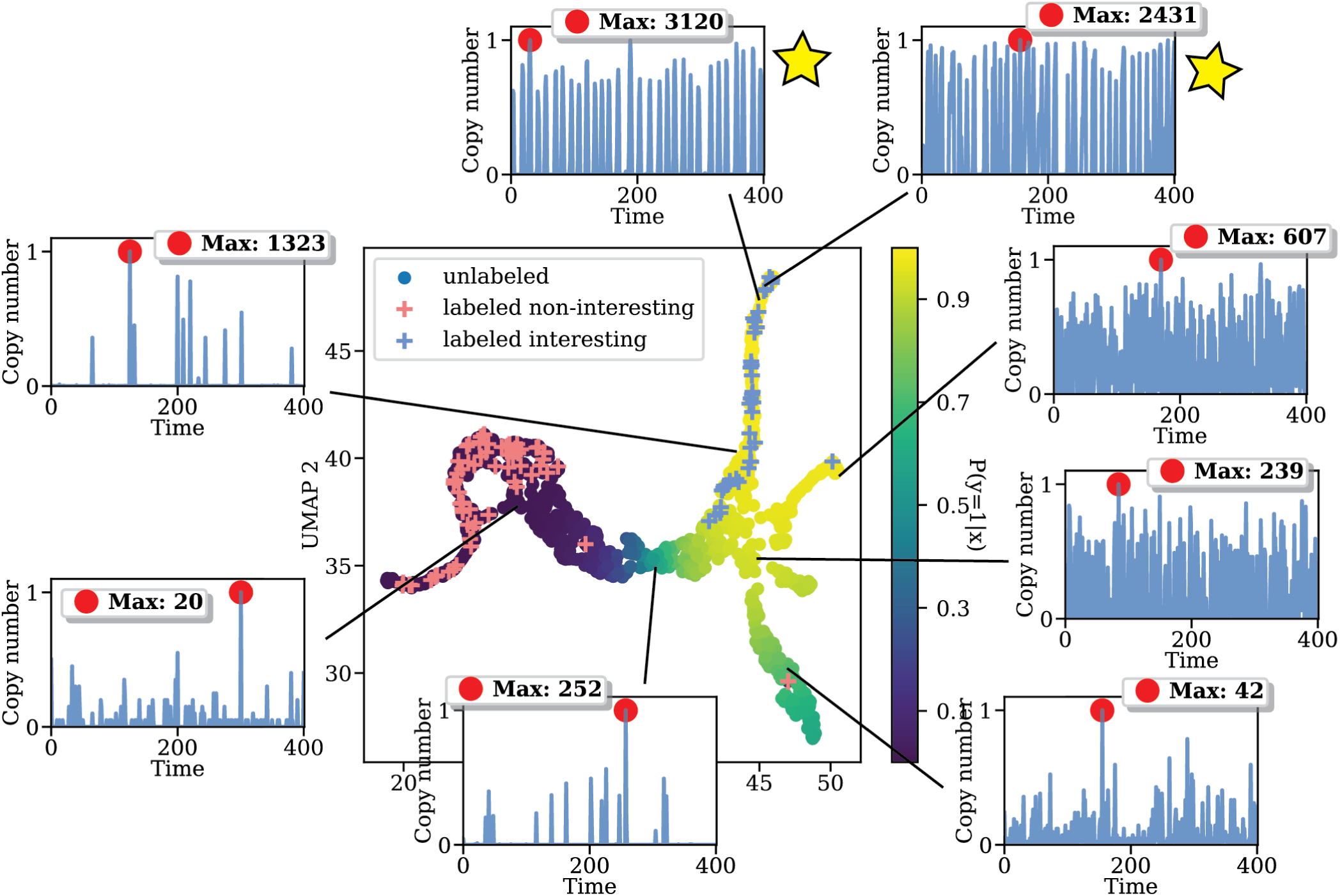
More samples added to the parameter sweep (947 points in total), a few more points has been labeled. Using label propagation gives predictions of unlabeled points. Yellow color of unlabeled points corresponds to a high probability of belonging to the interesting class (blue +, y = 1). Two trajectories of particular interest were found (star).

It is straightforward for the modeller to also map the gained label propagation information to parameter space. This can be used to fine tune the parameter sweep to particular ROIs. In our example, we were able to locate two trajectories of particular interests (Figure 2 star) which gave a high probability of belonging to the interesting class. Observe that the modeller is able to change their preferences at any time. This is one of the advantages of using a transductive semi-supervised approach such as label propagation. Once we found the trajectories in Figure 2 A and B, our preference was changed to focus on this particular ROI (Figure 3 A). Once the modeller have reached the point where she is satisfied with the current state of the system (i.e she have found some interesting trajectories and would like to find more of them) she can “freeze” the current state, i.e use the current dimension reduction mapping and labels to project new parameter points onto the same space and let the label propagation predict the outcome of these points. If new points get a low probability of belonging to the interesting class we will neglect them, thus the predictions from the label propagation will act as filter to filter out non-interesting realizations. To demonstrate this, we consciously sampled parameter points known to result in robust oscillations with a period of 24 hours and with copy numbers ranging between 1200-1500 for the Activator protein. Figure 3 B, show that all these parameter points become projected onto our ROI. Further, using a predictive probability threshold of 0.5, the label propagation will classify all of them as interesting. Note however, that the uncertainty within the ROI has increased. This can be due to that this behavior has not been seen before, i.e no labels has been associated with this behavior. Also note, that if we would have added new points that looked more similar to points labeled as interesting (i.e similar to Figure 2 A and B) these would probably get a higher probability, but using an uncertainty based measure such as entropy would have highlighted the robust oscillations.

**Figure 3.**
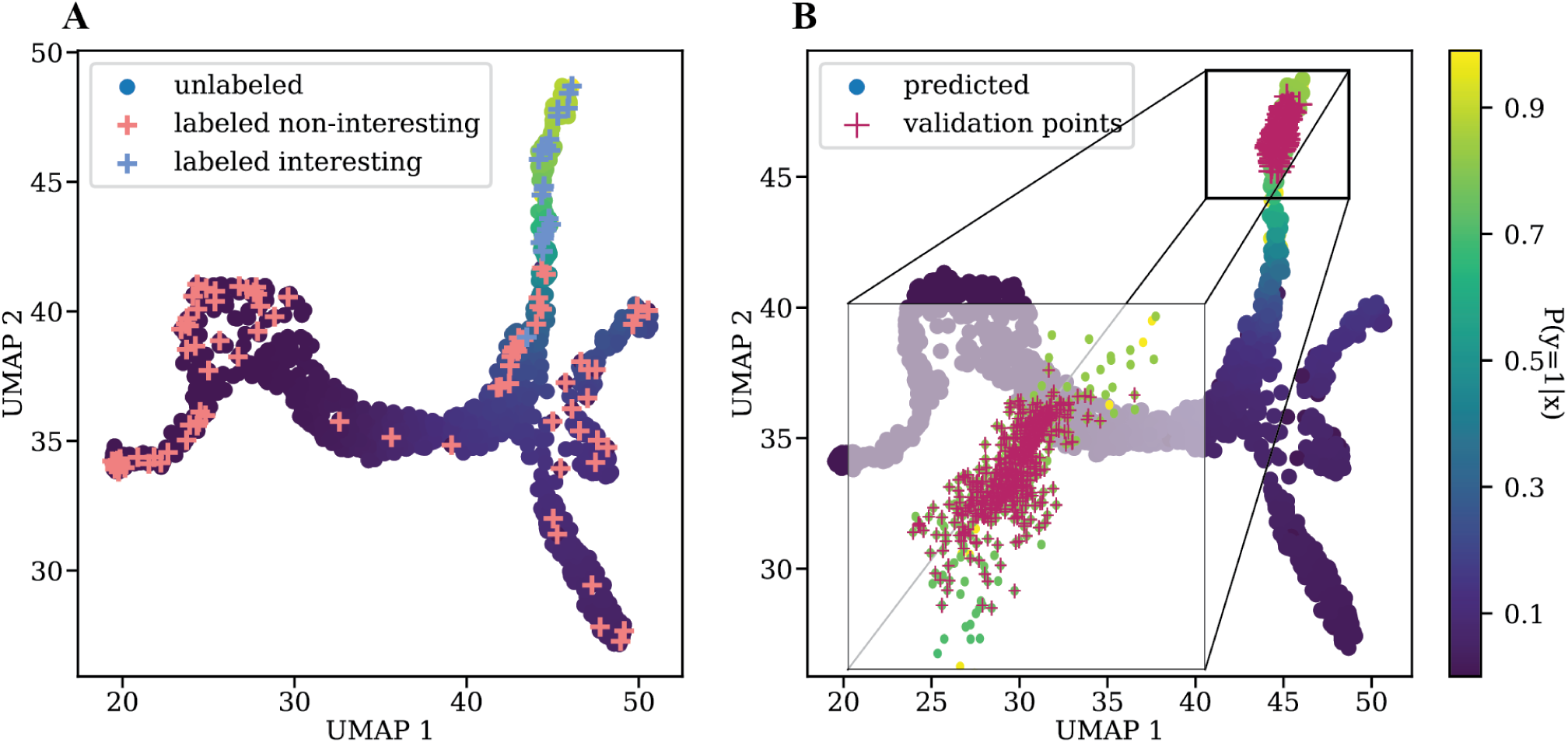
The parameter sweep continues. Here a total of 1553 samples are being observed. Label propagation enables the modeller to change and add labels to points as the sweep continues. Here, the focus and thus the preferences has been directed to the upper part of the data blob (A). A new label propagation is performed yielding new probabilities of unknowns. To simulate the process of a massive parameter sweep and the robustness of the current state of the system, we sample parameter points associated with robust oscillations. These points become mapped directly to our ROI (B), however the uncertainty grows, since this is a new and unseen behavior for the system. To reduce the uncertainty, the modeller can be queried to label points with high uncertainty using e.g entropy based measures.

#### Phase 3: Zooming in on the ROIs to engineer summary statistics

Phase 2 gave us a ROI which enables us to filter out non-interesting parameter points from the sweep using the dimension reduction mapping and a predictive threshold on the output from label propagation. Obviously, the modeller would like to inspect all points that pass the filter, i.e. “zoom in on the ROI”. To do this we simply create a separate projection using a DR method and explore the data (Figure 4 A). At this point in the smart workflow, it becomes more difficult to separate different behaviors within the ROI using only the initial minimal set of summary statistics. Ideally we could like to find new or additional summary statistics which can separate the robust oscillations from all other points in the ROI. To do this, we simply use our knowledge about suitable summary statistics to capture the oscillating properties of a time series, e.g features based on auto-correlation or Fast Fourier Transform (FFT). After some testing with different combinations of such summary statistics we found three which fitted well (Figure 4 B), namely 1. absolute sum of changes (Supplementary information), 2. mean aggregate of auto-correlation with various lags, and 3. variance aggregate of a auto-correlation with the same lags(Supplementary information). This process suggest that we can use sequences of ROIs to ultimately find more of the interesting trajectories and to engineer suitable summary statistics.

**Figure 4.**
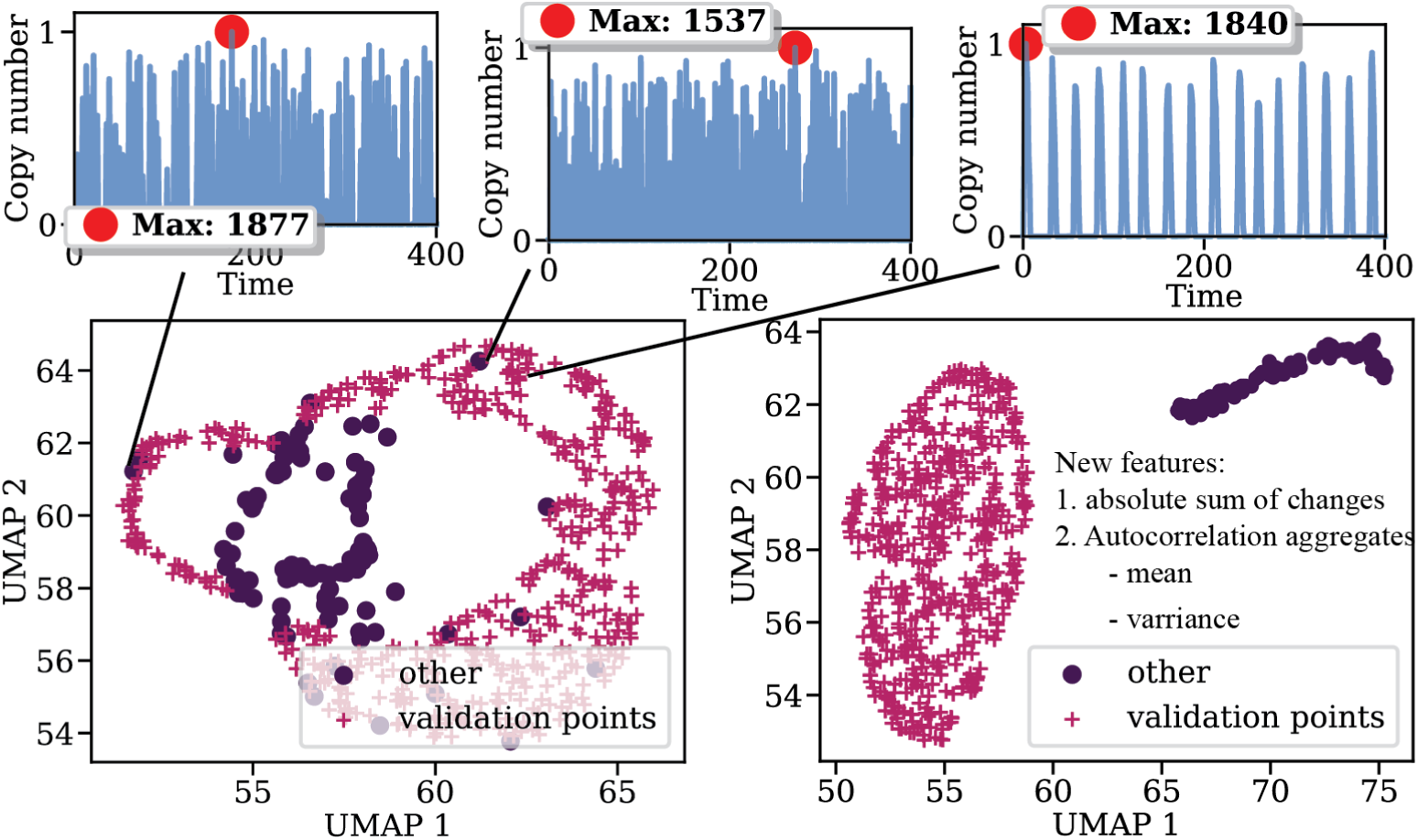
Zooming in on ROIs by neglecting points with low probability of beloning to the interesting class. We observe some non-robust outliers in the ROI (left). This suggest that we need other features than the minimal set to seperate these from the robust oscillations. By using features related to measures of oscillating patterns (e.g autocorrelation) we are able to separate the outliers in the ROI. This can be seen as a feature engineering for robust oscillations. Here we added “absolute sum of changes” and aggregates of autocorrelation features which clearly contribute to the separation from outliers (right).

## Discussion

Using techniques from semi-supervised machine learning, we have deviced a workflow for exploring high-dimensional stochastic models of gene regulatory networks that greatly reduce time to go from an initial prototype model to qualitative insights and predictions. Our human-in-the-loop exploration workflow effectively removes the need for hand-crafted analysis scripts based on prior information or hypotheses.

Using features to measure similarity between simulations results instead of the raw time series gives us several advantages. Mainly it opens up the opportunity for the smart system to be highly generalized to different models and objectives. The simulator itself can be regraded as a black-box and thus easily interchangeable. Further, it enables a wide range of machine learning techniques to be applied in the system, which do not have to be specialized for temporal data. Secondly, using features associated with time series analysis can be more informative and descriptive than raw time series and can further be used as summary statistics in complementary analysis such as likelihood-free parameter inference. The downside of using features is the computational burden of computing them. However, for the purpose of later large-scale downstream analysis of a model, a subset of features can be used. Note that, in our example we used our knowledge to test different summary statistics in a manual manner. However, in cases where this is not possible we can use feature selection techniques on a larger set of features. For example, the similarity matrix used in the label propagation is computed by the pairwise distance between points using the radial basis function (RBF) as a similarity measure. A feature selection approach could be to learn the weights (corresponding to the length-scale hyperparameter per summary statistics) for the RBF. Summary statistics with a small length scale would be given a higher weight for separating different labels. The learning could then involve optimizing e.g the mean label entropy the length scale by performing gradient decent^20^. An important note is to be cautious of overfitting when using a large set of summary statistics and performing feature selection, since we want our predictions to be generalized for new and unseen behaviors. This idea introduces us to another objective which has become apparent once we have used the system to label realizations. Our future objective is to build inductive classifiers which can predict newly generated realizations in a downstream process for massive parameter sweeps. Here, the classifier would act as filter which can structure the simulation results into qualitative interesting behaviors and e.g be used to save those realization into prioritized storage. This classifier would then be easily interchangeable between modelers and different models which contain similar qualitative behaviors. As an example, using the engineered features (Results: Phase 3) on the complete data set we are clearly able to linearly separate interesting points from non-interesting in the reduced dimensional space (Figure 5). This suggests that we can use a linear classifier as a compact model for filtering out robust oscillations in massive parameter sweeps. Moreover, a classifier like that could be used for more sophisticated and automatic approaches for sampling parameter points. By using sampling algorithms which utilizes both exploration (e.g space filling) and exploitation (i.e “zooming” in on ROIs or classified as interesting by a classifier) we would greatly reduced the effort needed to fine-tune parameter space searches. Such algorithms are often based on sequential Monte Carlo techniques^21^.

**Figure 5.**
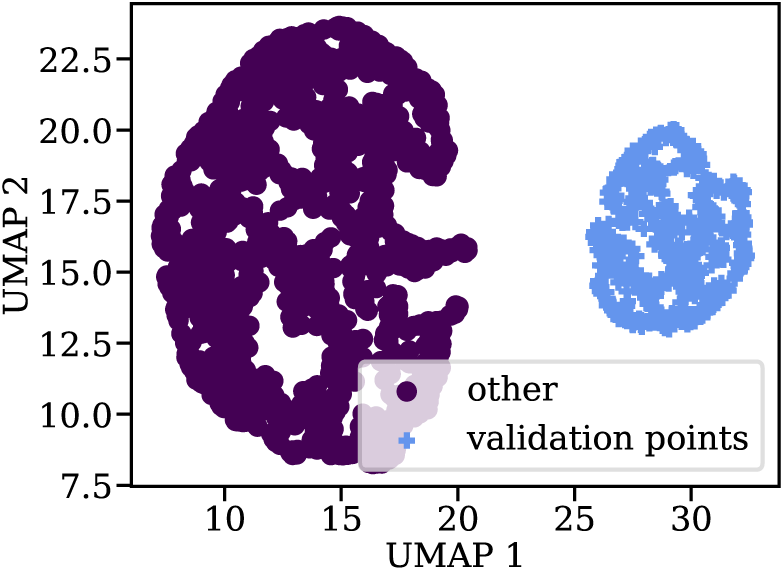
The complete sweep dataset with newly found features to separate noise from robust oscillations. Possible objective for classifiers, Monte Carlo estimation and likelihood-free parameter inference.

This work has been conducted in the context of the StochSS project^13^. StochSS, or Stochastic Simulation Service, is a cloud-native software as a service for constructing gene regulatory network models and to scale their simulations in public or private clouds^22^. In future work we plan to use the herein presented methodology to develop a Model Exploration Toolkit (MET); intelligent cloud services with the objective of letting users of StochSS rapidly input various versions of gene regulatory models and semi-automatically exploring them for interesting behavior. By training ensembles of inductive classifiers for different identified behaviors as outlined above, we have the opportunity to further automate identification of interesting dynamics by letting users share and deploy their classifiers. This vision outlines a collaborative modeling support systems that improves in the degree of automation by incorporating the knowledge of the expert modelers using it.

## Methods

### A smart workflow for model behavior discovery for high-dimensional models

The developed methodology is a smart workflow for screening and exploring a candidate model based on globally searching in the parameter space. Figure 6 outlines the basic approach: based on an initial sweep design, or placement of candidate points in parameter space, we generate simulation output data by running our simulator (which can be run in parallel). Time-series feature generation is conducted to generate summary statistics for each individual simulated parameter point. Based on these summary statistics, a dimensional reduction method projects the the large feature space into a two dimensional representation for visualization. The modeller is presented with an interactive 2D scatter plot of the reduced feature space where points in proximity to each other are assumed to have similar behavior, which can be observed by clicking on points showing a specified raw simulation output trajectory. At this point, the user can either choose to create a second sweep by modifying the initial sweep design, e.g to focus in on a few interesting parameter points to narrow down the search space. The second option (if sufficiently interesting points has been found) is to to give feedback about the most interesting parameter points by labeling them. Using this information, the system then proceeds to automatically label points which has not been labeled using a semi-supervised algorithm. This will enable the modeller to visually inspect points with high uncertainty, which in turn can be used to locate unseen or outlier points. Further, by using active learning (i.e letting the modeller label a few of the uncertain points) the system’s semi-supervised model will be updated and thus become more accurate. As the exploration of the data continues, the modeller is able to zoom in on particular interesting regions according to the labeling of the semi-supervised model, to enable an hierarchy of the exploration. This interactive human-in-the-loop machine learning approach is at the core of our workflow, and illustrated further in Supplementary Video 1. In the higher level exploration, the modeller will be able to tweak and engineer new summary statistics to fine-tune the separations of interesting and non-interesting trajectories. In the following paragraphs we will outline the specifics of each individual unit of the smart exploration workflow.

**Figure 6.**
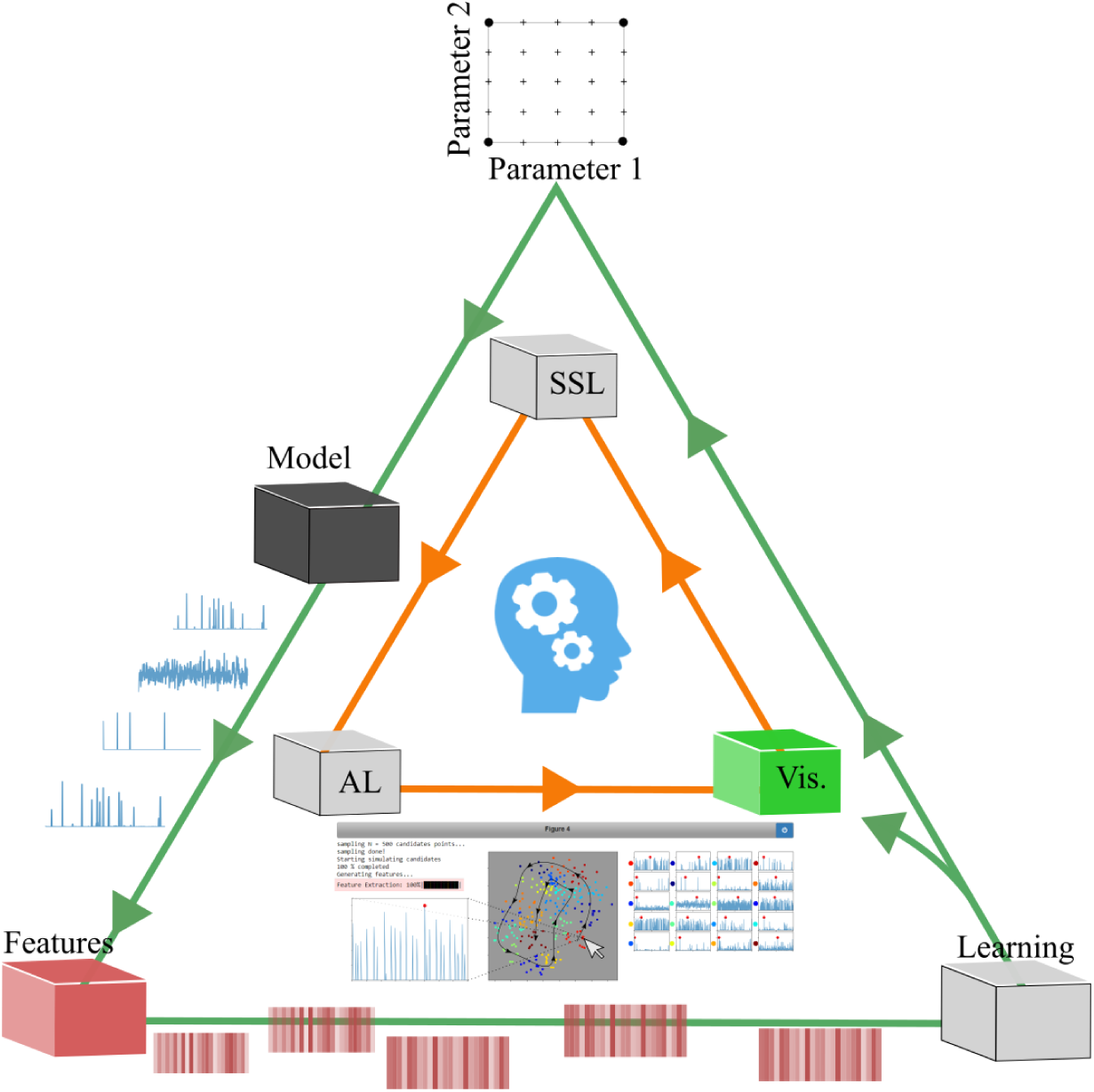
Smart Exploration workflow. The workflow starts with the user defining a parameter sweep design (top) to build a first batch of different parameter settings. Each parameter point is then realized in the model and corresponding simulator (black), which will output the trajectories for each realization. The raw trajectories (time series) are then converted to feature arrays (red) based on time series analysis. The data from the sweep will then enter a Learning cycle (white), where the modeler will be able to visualize the data points (associated with a realization) in a reduced 2D feature space and the corresponding trajectories (green). While exploring, the modeler can give feedback to the system by labeling data points. Only a few labeled data points are need since the data will then be fed into a semi-supervised learning algorithm (SSL). Using the semi-supervised model to infer labels of unlabeled data points, the system will also be able to evaluate the label uncertainty. This uncertainty can be used by active learning (AL) to ultimately suggest data points that the modeler might not yet discovered or to simply improve the semi-supervised model. Once the modeler is satisfied with the current labeling of data points the back-end workflow can be fine-tuned to include new parameter sweep designs.

### Parameter sweep design

To initiate the workflow we need an experimental design of the model parameter space. This involves defining global bounds for each parameter to define the search space in which parameter points (candidates) will be drawn under some probability distribution or picked in a deterministic manner. In case of little *a priori* knowledge of the model, one might for example start with a very simple exhaustive approach involving a uniform distribution bounded by minimum and maximum values for each parameter. A corresponding deterministic technique would be to sweep through a manually defined grid-search. Once a first batch has been processed by the workflow and we have gained more knowledge about the model behavior, these designs can be redefined according to the modelers preferences, e.g by mapping interesting realizations to parameter space and centralizing new designs around these corresponding parameter points. In the case of uniform sampling we have a parameter point

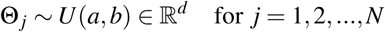

where *N* is the batch size and *a* and *b* are lower and upper bounds on the parameters. In practice, this range can be defined based on biophysical arguments, but it can be quite large (order of magnitudes).

### Stochastic simulation of gene regulatory networks

The most commonly employed mathematical framework for discrete stochastic simulations of chemical kinetics is continuous-time discrete-space Markov processes^23^, ^24^. The Stochastic Simulation Algorithm (SSA)^25^, commonly referred to as the Gillespie algorithm, generates statistically exact realizations from such processes. In our workflow the SSA will be treated as a black-box simulator, evaluated on a realization of the parameter points above

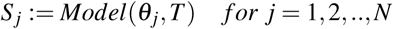

where *T* is the simulation time. Each realization of the model will contain the copy number evolution over *T* for all molecular species. We use the GillesPy^26^ package for simulations. GillesPy is part of the StochSS^13^ suite of tools, and wraps StochKit2^27^.

### High-throughput generation of summary statistics using comparative time series analysis

To represent simulation results in feature space rather then the raw simulation output, each batch of realizations passes a feature generating process where it is possible to generate several hundreds of features based on time series analysis. Each individual simulation result *S* _*j*_ corresponding to a particular realization of *θ*_*j*_ results in a feature vector

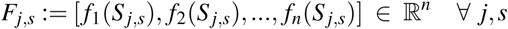

for a particular species *s* contains features *f* based on time series analysis, such as different moments, auto-correlation, Fast Fourier Transform (FFT) and many more. The system supports a minimal set of features to be generated for an initial large scale sweep design. The set of features can be manipulated to contain any feature set supported. We use the python package TSFRESH^28^(Time Series Feature extraction based on scalable hypothesis tests) for time series feature extraction. TSFRESH have a large library of time series feature functions, which ease the process of engineering features for various time series classification problems.

### Visualization and interactivity

Once the feature generation is finished the system will present the feature space in a reduced feature space using a specified dimension reduction (DR) method. Here, we use PCA^29^, kernel-based PCA^30^, t-SNE^17^ and UMAP^16^. While PCA is fast and enables inverse transform after projection, it does not capture non-linear relationships in the data. For this we either use t-SNE or UMAP. However, the support of several DR methods is useful for getting different perspectives of the data due to each method’s individual properties. Using interactive Jupyter notebooks^31^ as illustrated in Supplementary Video 1, the modeler can then navigate and explore different parameter points in a 2D scatter plot which is a mapping of the full feature space based on the dataset {*F*_*j,s*_}^*N*^ for a specified species *s*. By clicking on points, a specified trajectory of a particular species will show up next to the scatter plot. By using the assumption that data points in close proximity to each other have similar behavior, the effort needed to locate different or similar behaviors of the model is greatly reduced. The modeler will then have the opportunity to give feedback to the system about preferences over different model behaviors seen in the data by labeling individual points.

### Semi-supervised Label propagation and active learning

The labeling feedback can be used in a semi-supervised fashion for learning the preferred behaviors of unknown parameter points. This involves fitting a model to the data which models the associated labels (model behaviors) to parameter points which has not yet been looked at in the interactive exploration. If the model is probabilistic, we can infer some parameter points as being more uncertain than others. This enables us to: 1. direct the modelers attention to points that might not have been noticed by the modeler and 2. query the labels of these uncertain realizations to improve the accuracy of the model. The latter is commonly referred as active learning. Our current semi-supervised approach is label propagation by Gaussian Random Fields (GFR)^20^. In a semi-supervised setting, only a few points in the dataset are given a label, while the others remain unknown. GFR together with the properties of harmonic functions propagate the information of labeled instances to their neighbors. This follows from the assumption that nearby points (neighbors) have similar label information. By constructing a graph from the data, where nodes represent instances and edges the similarity between them, we can propagate the label information along the graph edges. The graph is represented by the *NxN* weight matrix *W* which measure the similarity between points. Using the Gaussian Radial Basis Function (RBF) as the similarity measure we get

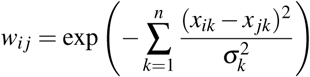

where *x ∈* ℝ^*n*^ can either be the full feature vector *F*_*j*_ belonging to parameter point *θ*_*j*_ for a specified species *s* or the same point represented in the reduced feature space (2D). *σ*_*k*_ is the length scale hyperparameter for feature *k*. We use the reduced dimensional space to propagate our labels, since this is the visual space that we explore during labeling. We also use the same length scale for both dimensions in this space. By partitioning *W* and the label vector *f* into partitions of labeled versus unlabeled instances, the label propagation over the unlabeled instances *y*_*u*_ corresponds to solving the linear system

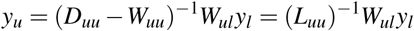

where *L* is the graph Laplacian and *D* is the diagonal matrix with entries *d*_*i*_ = ∑ _*j*_ *w*_*i j*_. We use the normalized graph Laplacian and remove clamping of labeled points, i.e labeled points can change their labels to some degree^18^. There is an elegant probabilistic interpretation of the label propagation above. Namely, it can be seen as a random walk procedure over the graph. The equation above can be expressed by the normalized transition matrix *P* = *D*^−1^*W.* Starting a random walk from an unlabeled node *i, P*_*i j*_ is the probability of walking (transitioning) to node *j* after one step. The walk will continue until a labeled node has been reached. *y*_*i*_ then becomes the probability of node *i* reaching a node of a particular label. The probabilistic framework enables the usage of active learning based on uncertainty of propagated labels over the graph. One simple way of measuring the uncertainty is by calculating the entropy

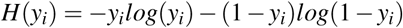

where a high entropy would correspond to high uncertainty. Thus, by arranging *H*(*y*_*u*_) we can query the modeler to label uncertain points, which will decrease uncertainty in the model and can also give new insight about the exploration.

### Hierarchical exploration by zooming in on ROIs to engineer summary statistics

At any stage after initiation of the workflow it is possible to “zoom in” on particular ROIs in the reduced feature space or in parameter space. Here we refer the “zoom in” as a way to isolate an ROI and explore it in more detail. This opens up the opportunity to fine-tune and engineer ad hoc summary statistics for the purpose of separating the interesting parameter points versus the non-interesting. This can either be done by manually adding new features using knowledge about suitable features, or to naively generate many features which then can be used for feature selection.

### Interactive parallel computing in clouds using Dask and MOLNs

The simulation (using SSA) and the feature generation are both embarrassingly parallel, and is well suited for parallelism in cloud computing infrastructure. We use the MOLNs^22^ orchestration software part of the StochSS suite of tools to deploy virtual clusters. For this work we have extended MOLNs with support for Dask, a framework for parallel computing in Python^32^. Apart from deploying Dask and Jupyter notebook, MOLNs installs the tools needed for scalable stochastic computational experiments of biochemical reaction networks. MOLNs is able to dynamically deploy clusters in a range of public and private clouds. This approach makes our software and workflow require minimal effort on setting up and configuring software, and allows for highly scaleable parallel computing. For the experiments in this paper we made use of the SNIC Science Cloud, a community cloud based on OpenStack, provided by the Swedish National Infrastructure for computing.

## Software

All code developed in this paper is available open source from: https://bitbucket.org/fwrede/psa

## Acknowledgements

The author’s are grateful for constructive discussions and feedback from Prashant Singh, Phil Harrison and Ola Spjuth. This work has been supported by the Uppsala Center for Interdisciplinary Mathematics graduate school, the Göran Gustafsson foundation and the NIH National Institute of Biomedical Imaging And Bioengineering (NIBIB) under award no. 2R01EB014877-04A1. All content is the responsibility of the authors and do not necessarily reflect the views of the funding agencies.

## Supplementary information

**Supplementary Video I.** This supplementary video illustrates an interactive model exploration using our tools, as described in the example Jupyter notebook (which is destributed with the sources code). Video avaiable here: https://www.dropbox.com/s/o0wszm7xdsnc7ri/paper1.mp4?dl=0

### Parameter sweep design of test GRN model

Table 1 outlines the parameter bounds used in the exploration of the oscillator and the robust parameter points for the validation points used in Results phase 3 and 4.

**Table 1.** Parameter Sweep design and robust settings for oscillations (24 h period).

| Parameter | lower bound | upper bound | robust |
| --- | --- | --- | --- |
| $\theta_A$ | 0.5 | 5000 | 50 |
| $\theta_R$ | 1.0 | 10000 | 100 |
| $\alpha_A$ | 0.5 | 5000 | 50 |
| $\alpha_A'$ | 5.0 | 50000 | 500 |
| $\alpha_R$ | 0.0001 | 1.0 | 0.01 |
| $\alpha_R'$ | 0.5 | 5000 | 50 |
| $\beta_A$ | 0.5 | 5000 | 50 |
| $\beta_R$ | 0.05 | 500 | 5.0 |
| $\delta_A$ | 0.01 | 100 | 1.0 |
| $\delta_{MA}$ | 0.1 | 1000 | 10 |
| $\delta_{MR}$ | 0.005 | 50 | 0.5 |
| $\delta_R$ | 0.002 | 20 | 0.2 |
| $\gamma_A$ | 0.01 | 100 | 1.0 |
| $\gamma_C$ | 0.02 | 200 | 2.0 |
| $\gamma_R$ | 0.01 | 100 | 1.0 |

### Summary statistics

Summary statistics (time series features from TSFRESH) used apart from the extreme values, mean, median, standard deviation, variance and sum.

- Absolute sum of changes:

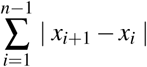
- Autocorrelation:

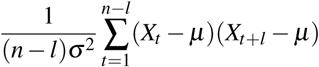

where *n* is the length of the time series *X*_*i*_, *σ*^2^ its variance and *µ* its mean. *l* denotes the lag.

